# Reducing Reactive Lipids Improves Cardiac Metabolic and Diastolic Function in Pulmonary Hypertension Models

**DOI:** 10.1101/2025.10.10.681763

**Authors:** James D West, Christy Moore, Santhi Gladson, Sheila Shay, Ethan Sevier, Elizabeth Kobeck, Ying Cai, Vineet Agrawal, John A Rathmacher, Anna R Hemnes

**Author notes:** Corresponding author: James D. West, Division of Allergy, Pulmonary and Critical Care Medicine, Vanderbilt University, 1161 21st Ave. S. Suite T-1218 MCN, Nashville, TN, U.S.A.

## Abstract

**Background:** Reactive oxygen species are increased across most pulmonary hypertension (PH) etiologies, resulting in increased reactive lipid dicarbonyls, which form protein adducts and impair mitochondrial function. We hypothesized that reducing reactive lipids would reduce right ventricular systolic pressure (RVSP) and improve cardiac function by eliminating protein-lipid damage feedback loops.

**Methods:** We used 2-hydroxybenzylamine (2-HOBA) to scavenge reactive lipids in three complimentary mouse models of PH: AKR-high fat diet (HFD, metabolic stress), LNAME-HFD (cardiometabolic syndrome), and pulmonary artery banding (PAB, load stress). Cardiac function was measured by echocardiography and catheterization. RV energy metabolism was determined by oxygraphy. Mass spectrometry analyzed lipids and ceramides; O-link and RNA-Seq evaluated proteomic and gene expression in lungs, RV, and LV.

**Results:** Reducing reactive lipids with 2-HOBA resulted in a ∼10% reduction in RVSP, reduced diastolic dysfunction, reduced plasma lipids and ceramides, and normalized RV fatty acid oxidation that was severely impaired in the AKR-HFD and PAB models. Proteomic and RNA changes in the lungs, RV, and LV suggested reduced oxidative damage and inflammatory signaling and altered developmental and actin organization signaling; these changes are plausibly associated with the improved adaptation. Some changes were sex specific, including a 4x higher cardiac fatty acid content in males than females.

**Conclusions:** Reactive lipid scavenging improves cardiac metabolic and diastolic function and pulmonary vascular resistance through restoration of mitochondrial function and reduced oxidative protein damage. The magnitude of hemodynamic improvement combined with substantial diastolic function improvement suggests clinical potential, particularly for PH patients with metabolic comorbidities.

## Introduction

Pulmonary Hypertension (PH) is a disease in which gradual cellular occlusion of the blood vessels in the lungs increases pulmonary vascular resistance, eventually leading to right heart failure and death. Although originating causes are different across patients, a common molecular factor across most etiologies is an increase in reactive oxygen species^3^ leading to(and originating from) a set of specific metabolic problems^1–6^.These include increased insulin resistance^2, 5, 6^, increased reliance on glutamine uptake in the lungs^1^, a shift away from normal TCA-glucose metabolism^4^, and suppressed fatty acid oxidation in the heart leading to increased intracellular fat^7, 8^. The severity of these metabolic disturbances may determine whether the right ventricle successfully adapts to increased afterload or progresses to failure^9–13^.

An important part of the mechanism is a feedback loop in which ROS causes reactive lipids particularly within the mitochondria, which then adduct to proteins causing loss of function and further metabolic defects (**Figure 1A**). A key target in PH is the mitochondrial lysine deacetylase, Sirt3, which is inactivated by lipid adduction^1, 4, 14, 15^. In a recent clinical trial attempting an intervention on this metabolic axis, the only patients in whom it succeeded were those with an unusual activating set of SNPs in Sirt3^4^, indicating that rescue of Sirt3 is a necessary prerequisite for resolving disease.

**Figure 1:**
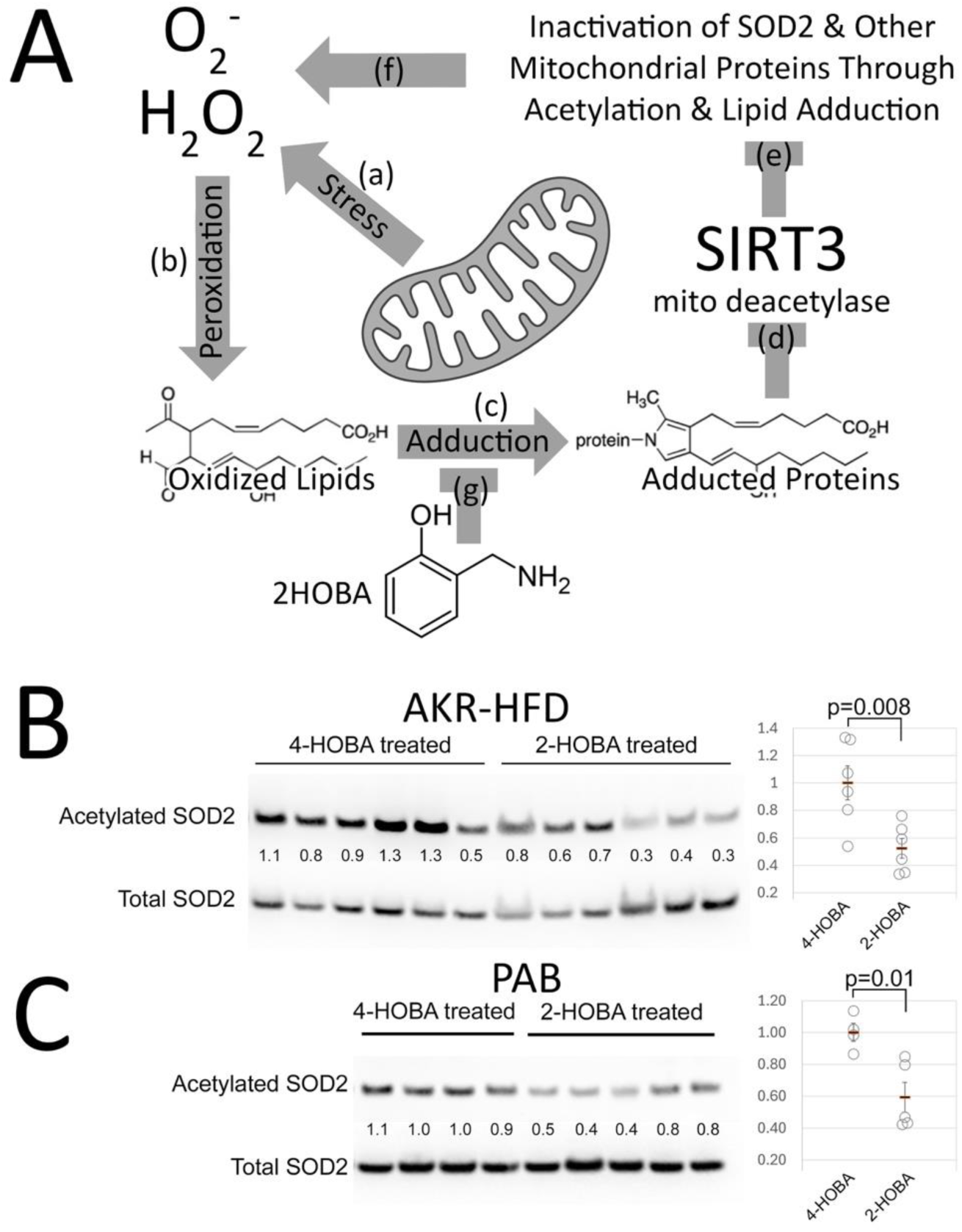
Proposed Mechanism. **(A)** When under stress (a), mitochondria produce more reactive oxygen species, including superoxide and peroxide, than the existing antioxidant pathways can handle. This rapidly results in creation of oxidized lipids (b) such as isoprostanes and isofurans. These reactive lipids adduct to proteins (c), generally reducing or altering protein function. One of these (d) is SIRT3, the mitochondrial deacetylase, which has reduced function as a result. One of SIRT3’s normal deacetylation targets is Superoxide Dismutase 2 (SOD2) (e), which is inactivated by acetylation. Loss of SOD2 and other mitochondrial protein functions results in worsened oxidative stress(f), in a vicious cycle. This can be interrupted (g) by scavenging reactive lipids using 2-HOBA. **(B-C)** In accordance with this mechanism, acetylated SOD2 is decreased by 2-HOBA in the AKR-HFD and PAB models respectively. Each lane is protein from one mouse; numbers are densitometry, plotted to the right. P value is unpaired t-test. LNAME-HFD is omitted because we had no available tissue after other assays.

This presents the possibility of rescuing the defects in metabolic function that are required for disease progression by alleviating the upstream problem with reactive lipid species. 2-Hydroxybenzylamine (2-HOBA) is a potent scavenger of reactive lipid dicarbonyls^16, 17^. It blocks the effects of reactive lipids by selectively trapping them – giving them an alternate target to bind to with a binding affinity two to three orders of magnitude stronger^16, 17^. We have published that 2-HOBA treatment rescued Sirt3 activity *in vivo* in BMPR2 mutant mice, resulting in normalization of metabolic markers and near normalization of pulmonary vascular resistance^1^, resulting in reduced acetylation of key targets. There’s thus strong rationale for intervention along this axis, and strong data that it can be effective, both mechanistically and physiologically. However, BMPR2 mutation is a relatively uncommon cause of PH, with very specific molecular defects downstream of the mutation. We don’t know whether 2-HOBA is similarly effective in other models. Further, although we know the immediate effect of 2-HOBA, we don’t know how this cascades through downstream molecular and physiologic mechanisms.

In the current study we tested reactive lipid scavenging in three complimentary mouse models representing distinct pathophysiologic mechanisms: AKR-high fat diet (metabolic stress) LNAME-HFD (cardiometabolic syndrome), and pulmonary artery banding (pure mechanical overload). This approach enabled therapeutic efficacy across different disease mechanisms while identifying universal effects from model-specific responses.

## Methods

All animal studies were approved by the Vanderbilt University Medical Center IACUC (protocols M2100005 (AKR), M2300008 (LNAME-HFD), M2200005 (PAB). Sample sizes were based on historical variation in primary outcome metrics (eg, RVSP, echo outcomes, O2K). For AKR-HFD and LNAME-HFD, all animals were included that survived to data collection. For PAB, our intent was to remove any animals with RVSP<35 (failed banding), but we had no animals that met this criteria. In each experiment, treatments were mixed within a day of assays to avoid confounding effects of day to day variation or mouse group.

### AKR-High Dat Diet Mouse Model

This strain develops metabolic syndrome and heart failure with preserved ejection fraction on HFD, though we acknowledge potential confounding from strain-specific pathologies including thyroid abnormalities and increased tumor susceptibility. Male AKR strain mice (n=16, 8-weeks old, Jackson Laboratories) were fed 60% fat diet (BioServ F3282, Flemington, NJ) and randomized to either 1g/liter 2-HOBA or 4-HOBA, an inert control (MTI Biotech, Ames, IA), in drinking water, with water changed weekly. Mice were weighed every two weeks. After 12 or 20 weeks, mice underwent echocardiography and closed-chested cardiac catheterization prior to sacrifice and tissue collection.

### LNAME-High Fat Diet Mouse Model

This model produces cardiometabolic syndrome with diastolic dysfunction while avoiding strain-specific confounders. C57BL/6J mice (n=32, 16 male/16 females, 8 weeks of age, Jackson Laboratories) received high fat diet (as above) and 0.5g/liter L-NAME plus 1g/liter 2-HOBA or 4-HOBA, as above, in drinking water, with pH adjusted to 7.2. Water was changed twice per week. Mice were weighed weekly. After 8 weeks, mice underwent echocardiography and closed-chested cardiac catheterization prior to sacrifice and tissue collection.

### Pulmonary Artery Banding Mouse Model

This model tests pure pressure overload effects independent of metabolic stress. C57BL/6J mice (n=32, 16 male/16 females, 8 weeks of age, Jackson Laboratories) underwent pulmonary artery banding (PAB) as follows. Mice were anesthetized with 2-3% isoflurane and orotracheally intubated. Animals were mechanically ventilated with isoflurane general anesthesia for PAB or sham surgery. A sternal incision was made using sterile technique, the pericardium was removed, and a partially occlusive titanium clip was placed around the pulmonary artery (Weck, Research Triangle Park, NC). The animal was then sutured closed and allowed to recover from anesthesia. Mice received 7.5 mg/kg carprofen by subcutaneous injection pre-operatively, and q12-24 hrs for 72 hours postoperatively. Sham treated animals underwent the same procedure without vascular clip placement. On post-op day 2, mice were randomized to 1g/liter 2-HOBA or 4-HOBA, as above, in drinking water. 3 weeks after surgery, mice underwent echocardiography and open-chested cardiac catheterization prior to sacrifice and tissue collection.

### Oxygraphy

The Oroboros O2K Oxygraph platform is used for precision OXPHOS analysis right ventricle heart tissue at controlled oxygen levels for high resolution respirometry. The right ventricle was harvested from mice and using a combination of mechanical separation and saponin permeabilization, and the tissue was prepped for analysis. Two to three milligrams of the tissue was then added to the O2K chambers that had been previously filled and calibrated with MiR05 respiration media. Once the chambers were sealed the mitochondrial function and oxygen flux per mg tissue in Fatty Acid Oxidation OXPHOS was assessed using a combination of substrate, uncoupler, and inhibitor reagents. The O2K’s high resolution polarographic oxygen sensor in conjunction with the DatLab software allows the user to obtain highly sensitive and quantitative real time oxygen concentrations and oxygen flux of the sample material. Personnel were blinded as to group identity during data collection. This data was then analyzed to assess differences in the mitochondrial function of Fatty Acid Oxidation in OXPHOS of 2-HOBA vs 4-HOBA treated Right Ventricle heart tissue.

### Hemodynamic Measurements

During all hemodynamic measurements, personnel were blinded as to group identity during data collection.

#### Echocardiography

Mice are anesthetized with 3% isoflurane on a heated pad in the supine position, depilatory cream applied to the thorax, and the mouse imaged with a Vevo F2. Mice are imaged along the parasternal long axis view, including a B-mode and 4D image; then a parasternal short axis view for an M-mode image across the papillary muscles. Next an apical 4 chamber view is taken for both pulse-wave (PW) Doppler imaging of LV blood flow, and imaging of the valves. Finally, the 4 chamber view were used for an M-mode cross section of the RV. All the images taken were loaded into the Vevo LAB software for analysis.

#### Closed-Chested Cardiac Catheterization

Mice were anesthetized with tribromoethanol (avertin), which does an excellent job of preserving heart rate. Right ventricular systolic pressure (RVSP) was measured by introducing a 1.4F pressure transducer into the right ventricle by threading through the right jugular, with the chest closed and the mice spontaneously breathing, as previously described^18^. Hemodynamics were continuously recorded with a Millar MPVS-300 unit coupled to a Powerlab 8-SP analog-to-digital converter acquired at 1000 Hz.

#### Open Chested Cardiac Catheterization

Open-chested catheterization was used for the PAB model because the clip interferes with the closed-chested method. Mice were anesthetized with 3% isoflurane and intubated with a 22 g catheter. Animals were mechanically ventilated at 18 cc/kg with vaporized isoflurane general anesthesia. Mice were positioned supine, ventral side up on a heated operating table at 37°C. A vertical incision over the abdomen was made and cautery was used to cut the diaphragm and expose the heart. A 1.4 French Mikro-tip catheter was directly inserted into the right ventricle. Hemodynamics were continuously recorded with a Millar MPVS-300 unit coupled to a Powerlab 8-SP analog-to-digital converter acquired at 1000 Hz.

### RNA-Seq

Lung, RV, and LV were flash frozen in liquid nitrogen immediately after sacrifice. RNA was isolated from whole lung using RNEasy kits (Qiagen), and delivered to Novogene (Sacramento, California) for paired end 150 sequencing on an Illumina platform. A nominal read depth of 20 million RNA (40 million ends) per animal was used. These were analyzed on the Partek platform, using the STAR aligner to align to the mm39 reference mouse genome. An average of 96% of reads aligned to genome. Counts were normalized to Counts Per Million. Group differences were assessed using ANOVA, and gene ontology determined using the R package clusterProfiler. Most statistical analyses and figure preparation were done within R. Data sets have been deposited in the GEO database at NCBI, as accession number (pending – GEO is closed for submissions during the shutdown).

### O-Link, Fatty Acid and Ceramide

Plasma, RV, and LV protein were submitted to the Vanderbilt University Medical Center High Throughput Biomarker Core for O-Link proteomics, and to the Bioanalytical * Drug Metabolism LC-MS Core for fatty acids and ceramides.

### Western Blots

Protein was run on a 4-12 % gel in MES buffer (NuPAGE) using antibodies to K68 actylated SOD2 (Abcam ab137037) and total SOD2 (Abcam ab13533).

### Statistical Analysis

Primary endpoints were analyzed using unpaired t-tests with significance set at p<0.05. RNA-seq differential expression required minimum 1.5-fold change and FDR<0.05. Sex differences were analyzed using two-way ANOVA when appropriate sample sizes permitted meaningful interpretation. Statistical tests were performed using the R stats package. Normality was established by Shapiro-Wilks test within R (RNA-seq data was log-transformed to achieve normality).

## Results

### Overview

This series of experiments was done sequentially over the course of two years, first AKR-HFD, then LNAME-HFD, then PAB. Thus, different measurements were taken as our understanding grew, explaining why the outcomes measured in each model aren’t identical. However, the data is more readily interpretable by presenting the same type of measurements across models, rather than by presenting them in chronological order.

We used three mouse models; in chronological order, these were. **(1)** The AKR-high fat diet (AKR-HFD) model was developed as a murine model of heart failure with preserved ejection fraction (HFpEF)^19^; AKR strain mice are genetically susceptible to PH with HFpEF-like features compared to other mouse strains. However, they have a variety of other health problems – in our studies, these included enlargement of the thyroid, fatty livers, and an assortment of tumors and leukemias. **(2)** The LNAME-HFD model consists of the combination of the nitric oxide synthase inhibitor Nω-nitrol-arginine methyl ester (L-NAME) with high fat diet for 8 weeks. This leads to a cardiometabolic syndrome related HFPEF, with generally mild to moderate LV diastolic dysfunction^20, 21^. **(3)** Pulmonary artery banding (PAB) was used to test a pure load stress model, in the absence of the high fat diet used in the other two models. A titanium clip is surgically placed around the pulmonary artery, resulting in significant increases in RV load. In all three models, 2-HOBA had the central molecular effects we expected, as assessed by reduced SOD2 acetylation (**Figure 1B-C**).

### 2-HOBA causes modest but significant reduction in RVSP, and improves diastolic dysfunction

2-HOBA reduced RVSP in both AKR-HFD and LNAME-HFD mice (**Figure 2A**), but not in the PAB model, in which the resistance is fixed by the band. The reductions were modest but statistically significant. Both cardiac output and ejection fraction were unchanged by 2-HOBA (**Figure 2B, C**), but neither were significantly reduced by the model, and so there was no decrease to correct. CO is much higher in the AKR-HFD mice – this may be a combination of larger size (they weighed nearly twice as much), different strain background, or a different echocardiography machine used for these. Fulton index scaled with RVSP as expected, but was not significantly affected by 2-HOBA (**Supplemental Figure 1A**), likely because the decrease in pressure is insufficient to see a change in muscularization given the variability and numbers. Mitral E wave – early (passive) filling of the ventricle – is lower in both LNAME-HFD and PAB models (not measured in the AKR), and is somewhat restored by 2-HOBA in the PAB, but not LNAME-HFD model (**Figure 2D**). Mitral A wave – late filling of the ventricle – is substantially reduced in the LNAME-HFD model, and almost entirely restored by 2-HOBA (**Figure 2E**). In the PAB model, we were hampered by limited effective measurements in the control (4-HOBA) mice, but still had significant restoration of the A wave velocity by 2-HOBA at every RVSP (**Figure 2E**). However, the A wave differences were only in males; in females, A wave appeared to be affected by neither the model nor treatment (**Supplemental Figure 1B**). Left atrial ejection fraction was also significantly higher with 2-HOBA in the LNAME-HFD model (**Supplemental Figure 1C**), supporting the A wave data. E/E’, which correlates to left ventricular filling pressure, is doubled in the LNAME-HFD model, and substantially improved by 2-HOBA (**Figure 2F**). E/E’ is normal in the PAB mice, and so unaffected by 2-HOBA (it can’t be improved beyond normal). Overall, these three metrics are strongly indicative of diastolic dysfunction, alleviated by 2-HOBA in both models, although with differences model to model.

**Figure 2:**
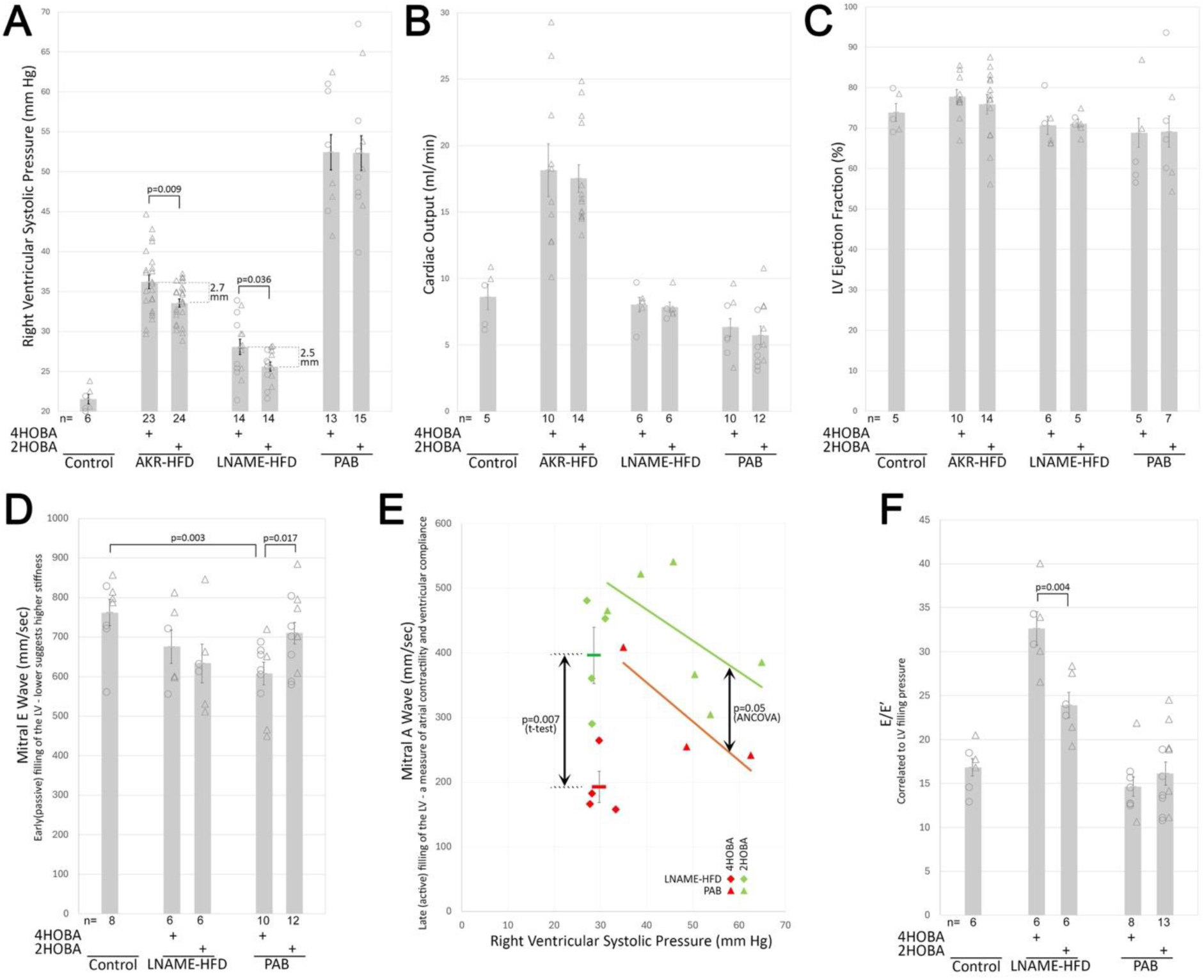
Hemodynamics. **(A)** 2-HOBA reduces RVSP in the AKR-HFD and LNAME-HFD models.**(B)** Cardiac output is relatively normal across models, and unchanged by 2-HOBA. **(C)** LV ejection fraction is relatively normal across models, and unchanged by 2-HOBA. **(D)** Mitral E Wave is reduced in both LHFD and PAB models, and rescued by 2-HOBA in the PAB model. **(E)** Mitral A Wave is reduced in both LHFD and PAB models, and rescued by 2-HOBA. **(F)** E/E’ is increased by LNAME-HFD, and rescued by 2-HOBA. Each symbol is a measurement from one mouse; circles are female; triangles are male (not different in this metric). For the AKR-HFD model, triangles to the left are 20 weeks of HFD; those to the right are 12 weeks (pooled because no differences were observed). Error bars are SEM. P-values are by t-test, except for ANCOVA in part E.

### Systemic effects include reductions in circulating LCFA and Ceramides, induction of signaling proteins, and reversal of high fat diet-induced neutrophilia

2-HOBA at least potentially has effects throughout the body, as does the high fat diet, LNAME, and at least potentially downstream effects of pulmonary artery banding. We thus were interested in impact on a variety of metrics in plasma, as well as metrics like weight gain and glucose.

We measured levels of 9 long chain fatty acids (LCFA) in plasma, including 14-0,15-0,16-0,17-0,18-0,18-1,18-2,20-0, and 22-0. Levels of four of these, included in **Figure 3A**, made up ∼97% of the total quantity, and were reduced 26% with 2-HOBA treatment in the AKR-HFD model, and 14% with 2-HOBA in the PAB model, which already had substantially lower circulating LCFA than the other models (**Figure 3A**), likely just because they did not have high fat diet.

**Figure 3:**
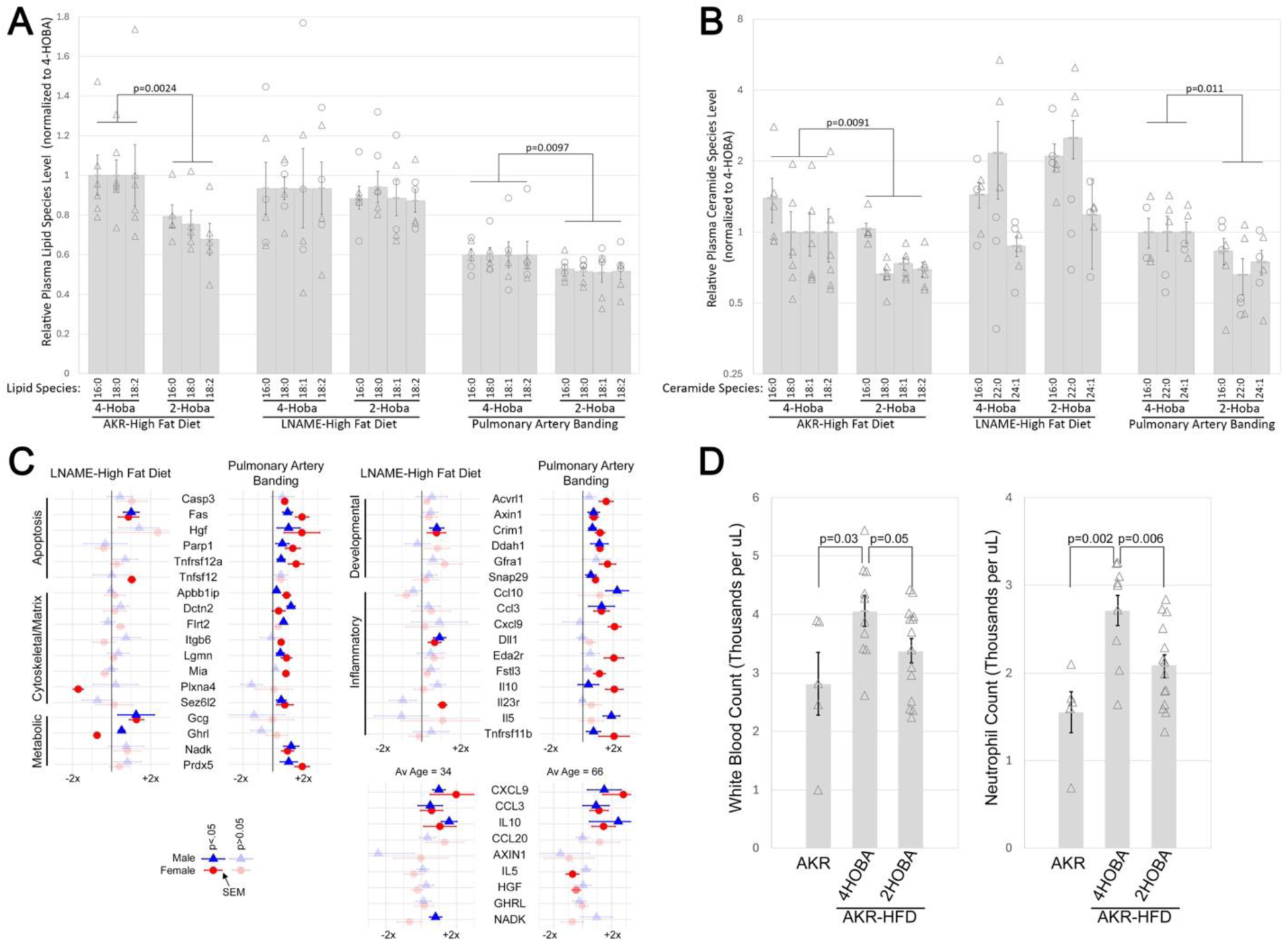
Circulating factors. **(A)** Plasma long chain fatty acids are reduced by 2-HOBA treatment in the AKR-HFD and PAB models, but not the LNAME-HFD model. **(B)** Circulating ceramides are reduced by 2-HOBA treatment in the AKR-HFD and PAB models, but not the LNAME-HFD model. **(C)** Multiple circulating proteins are altered by 2-HOBA treatment, although primarily in the PAB model, assessed by O-link. Symbols are average fold change comparing 2-HOBA to 4-HOBA, with error bars SEM. Triangles indicate males; circles females. 100% opacity indicates p<0.05 for different than 0 by unpaired t-test; low opacity are non-significant differences, included for comparison. **(C-lower)** similar data for 2-HOBA effect in two age cohort human patients by O-link somewhat parallels mouse data. **(D)** High fat diet induces increased neutrophil count in AKR mice, somewhat restored by 2-HOBA. Symbols as per A and B.

Ceramides are lipids that can be produced as intermediates of various synthesis or salvage pathways, but they’re also bioactive signaling molecules for stress responses, particularly oxidative or metabolic stress, although there is still lack of clarity around the meaning of different ceramides. Ceramides were measured in plasma from the three models; different ceramides measured in AKR than in the other two models, because of changes in the core protocol over time. 2-HOBA reduced average ceramide levels by 29% and 34% in AKR-HFD and PAB models respectively (**Figure 3B**), but not in the LNAME-HFD model.

We also performed O-link analysis for levels of 92 proteins in plasma from the LNAME-HFD and PAB models. 34 of these proteins were significantly changed in at least one of the models, for at least one sex (**Figure 3C**). Five of these were changed only in the LNAME-HFD model, two were changed in both models, and the other 27 were changed significantly only in the PAB model. Two of the genes unique to the LNAME-HFD model were glucagon (Gcg) and ghrelin (Ghrl), regulators of metabolism^22^. Nine of the proteins altered in the PAB model were also present in the O-link panel performed on plasma from human 2-HOBA trials^23, 24^. Three of these were also significantly changed across sexes and ages, and in the same direction as in mice, including the anti-inflammatory interleukin 10, and CXCL9 and CCL3.

We did not have O-link plasma data for the AKR-HFD model, but we did have CBC, which we don’t have for the other models. High fat diet is known to cause increased neutrophil counts^25^; this was true in the AKR-HFD model, but was substantially rescued by 2-HOBA treatment (**Figure 3D**).

2-HOBA did not affect weight, rate of weight gain, or blood sugar (**Supplemental Figure 2A, B**).

### Lung Effects of 2-HOBA in the AKR mice

We only analyzed the lungs closely in the AKR mice, since after that our focus shifted to the heart metrics. There were no significant differences between 2-HOBA and 4HOBA(control) treated groups in muscularized vessels or histology between AKR-HFD mice (not shown); the difference in RVSP was low enough that it wouldn’t be expected.

We performed bulk RNA-seq on these lungs. There were 325 genes differentially regulated at a minimum of 1.2 fold change and p<0.05 (raw t-test) (**Supplemental Table 1A**). 84 of these have no gene ontology information; of the remaining 241, 101 (42%) map to overrepresented gene ontology groups (**Supplemental Table 1B**). 76 of these are represented in the 17 GO groups in **Figure 4A**.

**Figure 4:**
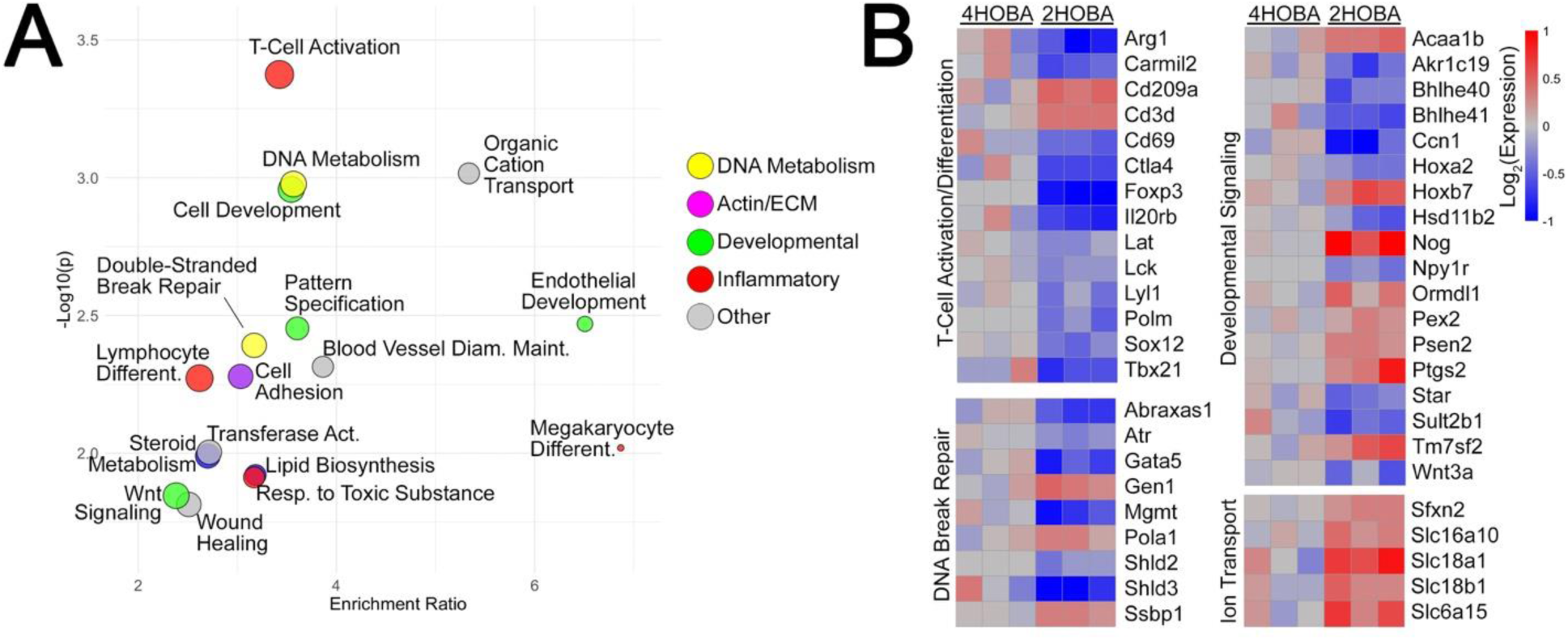
Lung Gene Expression Changes. **(A)** Gene ontology groups of significantly altered genes in 2-HOBA vs 4HOBA treatment by bulk RNA-seq of lung in AKR-HFD mice. All were done in male mice, with three mice in each treatment group sequenced. Dot size and is proportional to number of genes in the group; color is by rough grouping of ontology categories. **(B)** Heatmap of representative genes from these categories, normalized to average of 4HOBA treated mice. Each block is gene expression from one animal.

Selected genes from these are plotted in **Figure 4B**. Most T-Cell activation genes are decreased, but Cd3d is increased, suggesting that the overall level of T-cells likely stays the same or increases – but the activation levels decrease.

There’s an increase in a variety of ion transport genes – but these are actually expressed in different cell types; based on published single cell sets, for instance, Sfxn2, Slc18a1, and Slc6a15 are primarily expressed in various epithelial types, while Slc16a10 is primarily expressed in monocyte-derived macrophages (MDM) – the increase there is comparable to the increase in Cd209a, an MDM marker.

There is a broad decrease in most DNA repair genes, which makes sense since reactive lipids cause significant DNA damage, but a few specific pathways are increased – for instance, Gen1, which resolves Holliday junctions^26^, is increased.

The same is true for the developmental genes – a mixture of up and downregulated genes, complicated by different locations of expression. Nog is strongly upregulated, and is primarily expressed in capillary endothelium; Wnt3a is downregulated, and primarily expressed in AT1 epithelium; Npy1r is downregulated, and primarily expressed in fibroblasts.

Overall, the most clearly interpretable changes in the lungs caused by reduction in reactive lipids are associated with reduced T-cell activation and likely reduced DNA damage. These are plausible mechanisms^27–29^ for the reductions in pulmonary vascular resistance we saw.

### 2-HOBA improves impaired RV energy metabolism, but with possible sex differences

We measured metabolic activity in the right hearts of mice, using the Oroboros O2k. RV were removed from mice immediately after sacrifice and oxygen consumption in response to different substrates tested. We found that, generally, where oxygen consumption was impaired by the model, 2-HOBA at least partially restored it.

In the AKR model, we only used males, but oxygen consumption with 2-HOBA was significantly improved across substrates, although likely driven by restoration of impaired fatty acid oxidation (**Figure 5A**). The LNAME-HFD model had essentially no impairment of oxygen consumption in males, and so there was nothing for 2-HOBA to restore. In the females, there was mild impairment, which was statistically significantly restored to normal by 2-HOBA (**Figure 5B**). In the PAB model, males had significantly reduced oxygen consumption, substantially restored by 2-HOBA. Females had more moderate reductions, but 2-HOBA did not significantly improve it (**Figure 5C**).

**Figure 5:**
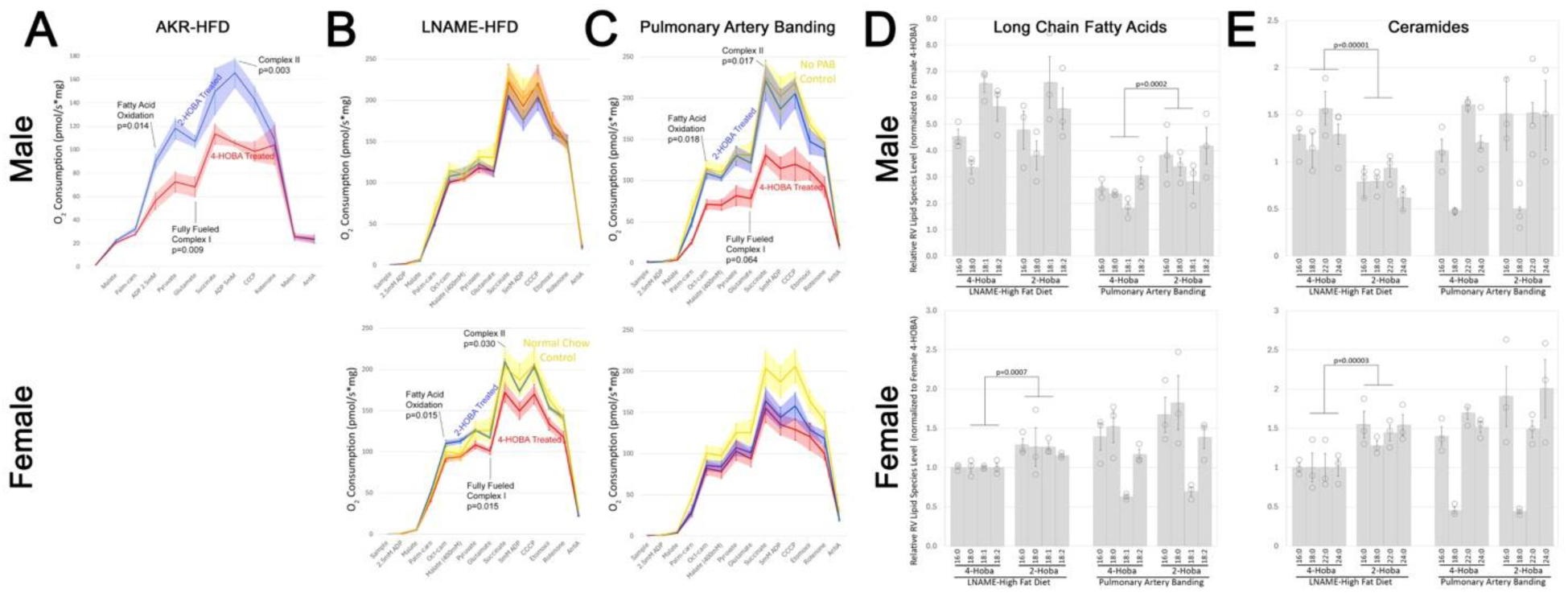
RV Energy metabolism, by sex. **(A-C)** 2-HOBA restores impaired oxygen consumption by substrate in fresh RV from male (top line) or female (bottom line) mice, assessed by O2k. Lines are the averages with shading at the level of error bars, which are SEM. Yellow lines are strain-matched no treatment controls; red lines are 4-HOBA treated; blue lines are 2-HOBA treated. **(D)** Long chain fatty acid species, determined by mass-spec. Each symbol is measurement from one animal; columns are averages, error bars are SEM. All values are normalized to female 4-HOBA LNAME-HFD values, to improve readability. Statistical values are unpaired t-test. **(E)** Ceramide species, determined by mass-spec, as for (D).

We measured long chain fatty acids (LCFA) in the RV for the LNAME-HFD and PAB models. As for plasma, more than 97% of the LCFA were included in 16:0, 18:0, 18:1, and 18:2. We normalized to female 4-Hoba in LNAME-HFD to make it easy to read (this doesn’t change outcomes from non-normalized). First, in both LNAME-HFD and PAB, males had significantly higher RV LCFA of every species than females. Females, but not males, had a statistically significant increase in LCFA with 2-HOBA in the LNAME-HFD model, while males, but not females, had a statistically significant increase with 2-HOBA in LCFA in the PAB model (**Figure 5D**). This may be coincidence, but these are the same sex-model pairs in which 2-HOBA improved energy metabolism; increasing LCFA was thus associated with restoration of impaired energy metabolism.

Finally, we measured ceramide levels in the RV. Ceramides weren’t affected by 2-HOBA treatment in the PAB model, but went in opposite directions by sex in the LNAME-HFD model; they were significantly decreased in males, but significantly increased in females (**Figure 5E**). Since this increase in females was associated with improved energy function (**Figure 5B**) and reduced diastolic dysfunction (**Figure 2**), interpretation isn’t immediately obvious; ceramides are generally thought to be harmful, but increased levels of C22 and C24 have been associated with reduced risk of heart failure^30^.

### Expression and Protein Expression Changes in the RV with 2-HOBA in the three models

We examined bulk RNA-seq from the RV in the three models; in addition, we were able to collect O-link data on the LNAME-HFD and PAB models. These were only done in male mice, primarily for consistency, although it does mean that we’re unable to use this to probe the sex differences seen above. The three models had substantially different specific genes altered, which is unsurprising given the substantially different stresses and strain differences.

In the AKR right ventricles, there were 152 genes altered greater than 1.5x, at p<0.05, by 2-HOBA treatment. These fell into gene ontology groups with clear relevance to the phenotype described above, including regulation of inflammatory response, metabolic substrate transport, and muscle differentiation, among others (**Figure 6A**). In the LNAME-HFD ventricles, there were 168 genes altered greater than 1.5x, at p<0.05, by 2-HOBA treatment. The groups altered were for the most part different than those in the AKR mice, with the exception of ERBB signaling, and were more dominated by metabolic ontology groups (insulin, glucose transport, mitochondrial organization) and a different set of signaling pathways(**Figure 6B**). These still have clear relevance to the phenotype. In the PAB ventricles, there were 159 genes altered greater than 1.5x on average, at p<0.05, by 2-HOBA treatment. As one might expect from lack of a direct metabolic stimulus, none of the overrepresented gene ontology groups are directly metabolic in nature, but are instead related to inflammation, proliferation, and adhesion. (**Figure 6C**). Note that slightly different choices of minimum p value or fold change makes differences to these numbers, but overall the dominant gene ontology groups remain the same.

**Figure 6:**
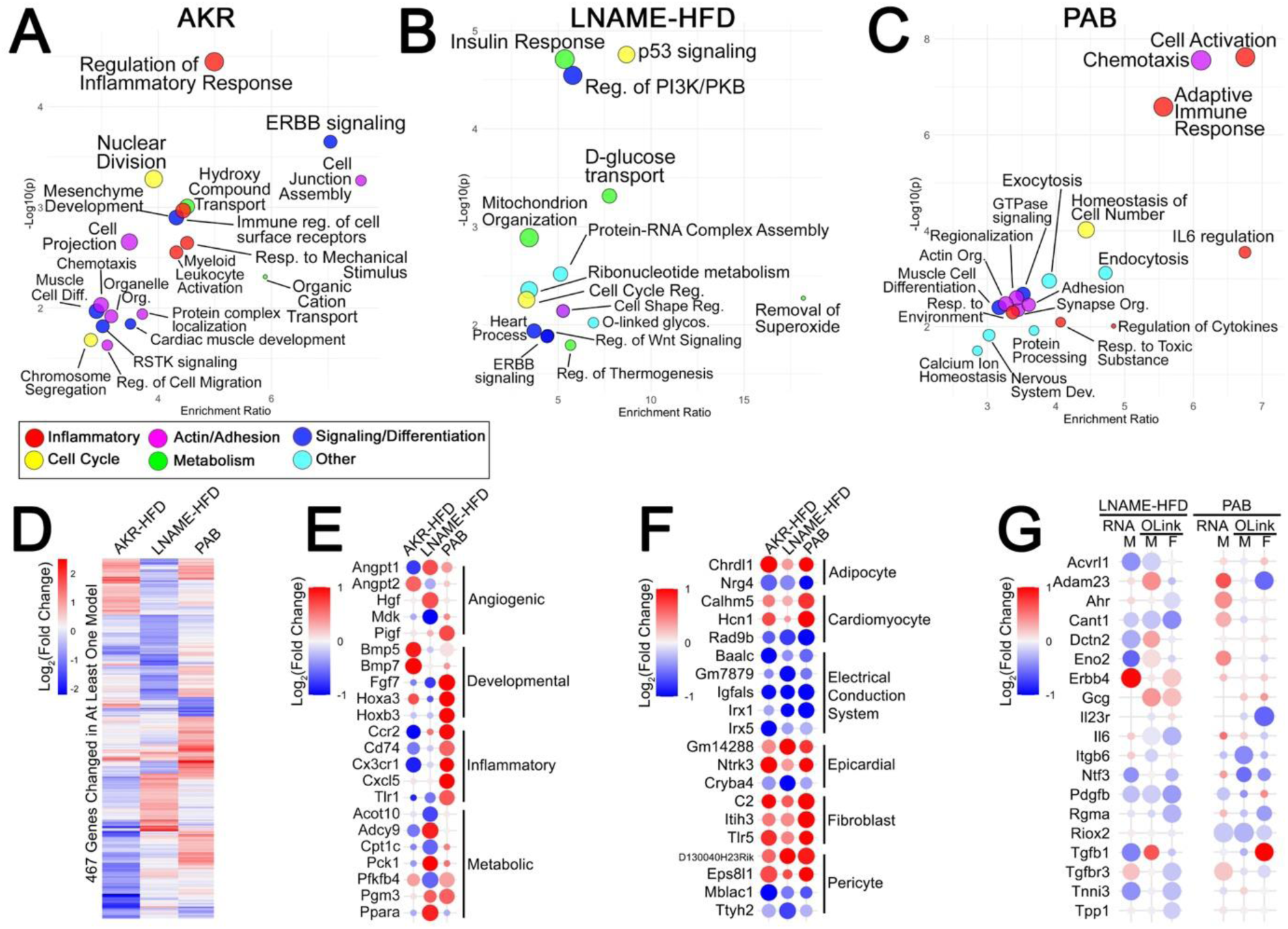
RV Molecular Changes. Gene ontology groups of significantly altered genes in 2-HOBA vs 4HOBA treatment by bulk RNA-seq of right ventricle (RV) in **(A)** AKR-HFD **(B)** LNAME-HFD and **(C)** PAB mouse models. All were done in male mice, with three mice in each treatment group sequenced. Dot size is proportional to number of genes in the group; color is by rough type of group. **(D)** heatmap of average fold change between 4HOBA and 2-HOBA treated mice, for all 467 genes changed in at least one experiment, shows relatively little overlap in changes across all three experiments. Red genes are increased with 2-HOBA; blue are decreased. **(E)** Examples of genes from different ontology categories across models. **(F)** Some genes are changed in common across experiments, but many of these are primarily expressed outside of cardiomyocytes. **(G)** Significant protein changes by O-link, with RNA-seq results for comparison. RNA-seq results are all male; some of the changes in protein are sex-specific. **For (E,F,G)** - Color indicates fold change; size indicates significance – sizes below 80% of maximum are p>0.05, so many of these are merely trending.

Taken together, these are 467 genes – and there’s relatively little congruence in which genes are changed across models. The heatmap in **Figure 6D** is illustrative – although some gene changes may be in common across two models, very few are in common across all three. These findings support the hypothesis that reactive dicarbonyl lipids are a common upstream mediator of diverse downstream pathways in these models, indicative of a common nodal pathway that may give rise to the heterogeneous phenotypes in Group 2 PH. In other words, we’d expect different models to have different impacts of reactive lipids.

It’s worth noting that while the specific genes are different, likely driven by differences in the models, in general there are angiogenic, developmental, inflammatory, and metabolic changes across models (**Figure 6E**). It’s also difficult to interpret up vs down-regulated in gene expression – it depends on whether gene expression is driving, or responsive to, changes in protein or function.

There are, however, some genes that are changed concordantly across experiment types. Although this was bulk RNA-seq, not single cell, many of the genes consistently changed across experiments are primarily expressed in supportive cell types, not cardiomyocytes (**Figure 6F**). Some of these changes have existing publications suggesting they’re cardioprotective^31–33^. However, because most genes in these minority populations would be hidden in the bulk RNA-seq, it’s difficult to tell exactly what’s happening. Overall, the bulk RNA-seq shows that the response is distinct to the model system – but that beyond that, it’s difficult to understand details without single cell RNA-seq, which we don’t have currently.

We also used the mouse O-link panel to examine levels of 92 proteins in the RV in the LNAME-HFD and PAB models. 19 of these were differentially regulated between 4HOBA (control) and 2-HOBA treated animals in at least one of the experiments (**Figure 6G**). Comparing the protein data to the bulk RNA-seq data continues our story of complexity. In 4 of these genes, RNA-seq matches protein data. For Acvrl1, Cant1, Pdgfb, Riox2, direction of protein change in males (or males and females) matches the direction of RNA change. For 3, the data matches, but in the wrong sex – Erbb4, Ntf3, Tnni3 have changes in protein in females, but not males – but changes in RNA in the males. For 5, the changes are discordant – Adam23, Dctn3, Eno2, Tgfb1, and Tgfbr3 have directions of protein change in the opposite of RNA change. For 2, Gcg and Il23, RNA levels were below detection. And for the remaining 5, Ahr, Il6, Itgb6, Rgma, and Tpp1, changes in protein are associated with no change in RNA levels. In interpreting this, one must recall that 2-HOBA is not targeting a specific receptor; it is preventing lipid adduction to random proteins caused by oxidative stress and lipid oxidation, which maintains protein function, and the feedback to gene expression is thus extremely indirect and moderated by functional changes.

### Expression, Protein and Fatty Acid Changes in the LV with 2-HOBA in the three models

The AKR-HFD and LNAME-HFD models are models of heart failure with preserved ejection fraction, characterized by diastolic dysfunction, as seen in **Figure 2**. We were thus also interested in the effects of 2-HOBA on the left ventricle (LV).

We measured long chain fatty acids (LCFA) and ceramides in the LV by mass-spec in the LNAME-HFD and PAB models. As for RV and plasma, we focused on the LCFA which make up the majority of the species. The first striking finding is that, unlike the RV, in the LV there is not a difference in LCFA levels between males and females (Compare **Figure 5D** to **Figure 7A**). Similarly, there’s not a difference in 2-HOBA effects between males and females; in both models, there’s a slight increase in 3 of 4 of the dominant LCFA. Ceramides had a similar distribution as was found in the RV (**Figure 7B**), but although there was a trend toward change in some of the species, it was not affected by 2-HOBA in either model.

**Figure 7:**
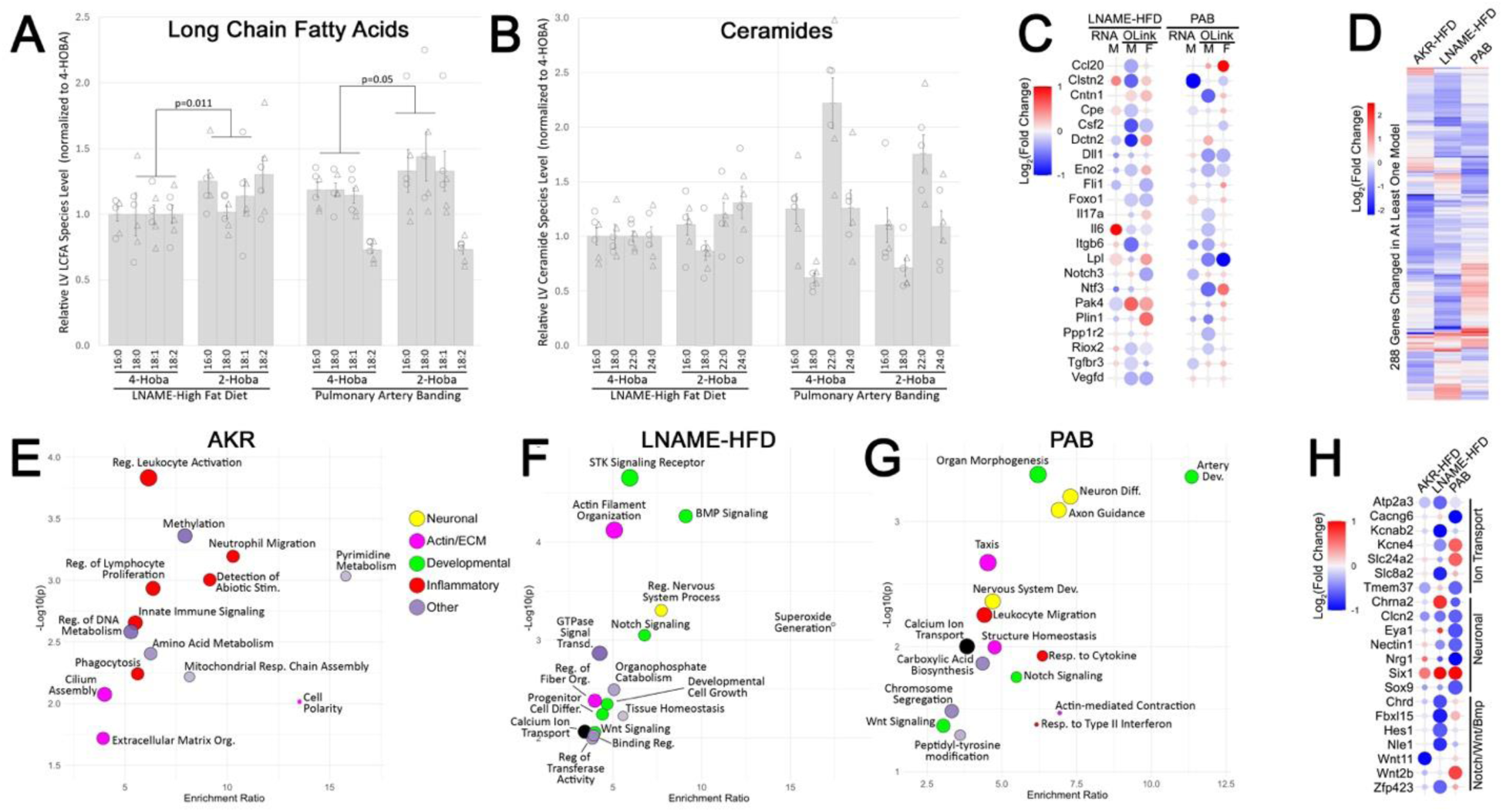
LV Molecular Changes. **(A)** Long chain fatty acid species, determined by mass-spec. Each symbol is measurement from one animal; circles are female, triangles male, which were not different in the LV. Columns are averages, error bars are SEM. All values are normalized to 4-HOBA LNAME-HFD values, to improve readability. Statistical values are unpaired t-test. **(B)** Ceramide species, determined by mass-spec, as for (A). **(C)** Significant protein changes by O-link, with RNA-seq results for comparison. RNA-seq results are all male; some of the changes in protein are sex-specific. Color indicates fold change; size indicates significance. **(D)** heatmap of average fold change between 4HOBA and 2-HOBA treated mice, for all 288 genes changed in at least one experiment. Red genes are increased with 2-HOBA; blue are decreased. **(E, F, G)** Gene ontology groups of significantly altered genes in 2-HOBA vs 4HOBA treatment by bulk RNA-seq of left ventricle (RV) in **(E)** AKR-HFD **(F)** LNAME-HFD and **(G)** PAB mouse models. All were done in male mice, with three mice in each treatment group sequenced. Dot size is proportional to number of genes in the group; color is related to the type of ontology group. **(H)** Examples of genes from different ontology categories across models.

As was done for RV, we measured 92 proteins with O-link in LV tissue from the LNAME-HFD and PAB models. 22 of these were differentially regulated between 4HOBA (control) and 2-HOBA treated animals in at least one of the experiments (**Figure 7C**). 7 of the 22 are in common with proteins changed in the RV, and in 5/7 the change was concordant with the RV change. As was true for the RV, bulk gene expression does not generally match the protein changes. The proteins changed have a variety of functions.

Using the same criteria for looking at bulk RNA-seq data as was used for RV, there were 97 genes altered greater than 1.5x, at p<0.05, by 2-HOBA treatment in the AKR-HFD model, 115 in the LNAME-HFD model, and 83 in the PAB model. There are only 7 genes shared between lists, for a total of 288 genes changed by these criteria in at least one model. Only 30 (10%) have concordant changes of >1.2x in all three model, but 136 (47%) are down at least 1.2x in 2/3 models, and another 19 (7%) are up at least 1.2x in 2/3 models(**Figure 7D** shows average fold change across the models)

Considering the gene ontology groups these genes fall into, AKR is dominated by inflammation-related groups, with some matrix/motility related, and some energy and anapleurotic metabolism (**Figure 7E**). LNAME-HFD is quite different, with no inflammation groups, but a lot of developmental and signaling groups, with prominent actin organization (**Figure 7F**). PAB is more similar to LNAME-HFD, with developmental and actin groups, but it has very prominent groups related to neuronal development (**Figure 7G**). Both LNAME-HFD and PAB have alteration in ion transport genes, some of which are also plotted in **Figure 7H**. The genes included in the neuronal category include many which are only neuronal, implying changes in one of the electrical conduction cell types in the heart, rather than in cardiomyocytes or vascular cells.

The key points here are that the left ventricle is also substantially affected by these models; that reduction in reactive lipids with 2-HOBA effects changes on the level of lipids, proteins, and gene expression that are largely but not entirely distinct to the models; and that the changes likely include many cell types beyond cardiomyocytes or fibroblasts.

## Discussion

Group 2 PH remains the most common cause of pulmonary hypertension worldwide, independently increases morbidity and mortality when identified in left heart disease, and yet currently has not directed therapies to improve mortality. A significant challenge in identifying therapeutic targets has been the heterogeneous nature of Group 2 PH. Despite the heterogeneity in Group 2 PH, there may be common upstream or downstream pathways that drive progression of the disease, regardless of the initial cause, but no studies have yet identified these common pathways. Our findings support that reactive dicarbonyl lipids may represent a common pathway that drives a component of Group 2 PH regardless of the initial cause, and similarly to therapies such as SGLT2 inhibitors that demonstrate ubiquitous benefit in heart failure regardless of cause, targeting reactive dicarbonyl lipid species therapeutically may also be ubiquitously efficacious.

This study demonstrates that reactive lipid scavenging with 2-HOBA provides therapeutic benefit in preclinical models of pulmonary hypertension. Three lines of evidence support our hypothesis that 2-HOBA prevents oxidative stress-mediated protein damage through non-specific prevention of lipid-protein adduction rather than by targeting particular pathways.

First, 2-HOBA produced consistent physiological improvements across all three mechanistically distinct PH models. We observed 8-12% reductions in pulmonary vascular resistance (**Figure 2A**), improvements in diastolic function markers (**Figure 2D-F**), and restoration of right ventricular fatty acid metabolism (**Figure 5A-C**). The consistency of these beneficial effects despite different disease mechanisms supports a system-level intervention acting on oxidative damage rather than model-specific pathway inhibition.

Second, the molecular responses to 2-HOBA were markedly model-specific even as physiological improvements remained consistent. Gene ontology analysis revealed distinct dominant signatures: reduced inflammation in the AKR-HFD model, altered energy metabolism in the LNAME-HFD model, and immune and cytoskeletal changes in the PAB model (**Figure 6**). This molecular heterogeneity accompanying consistent functional benefit strongly supports the hypothesis that 2-HOBA acts by preventing random protein adduction across diverse cellular contexts, rather than modulating specific signaling pathways. The proteins affected appear to be those most vulnerable to lipid adduction in each disease setting.

Third, we observed broad improvements across multiple biological levels—plasma lipids and ceramides were reduced in both AKR and PAB models (**Figure 3A-B**), lung gene expression showed reduced T-cell activation and DNA damage responses (**Figure 4**), and right ventricular metabolic capacity was restored. This multi-system coordination suggests that reducing the burden of non-specific protein damage allows normal regulatory mechanisms to reassert control, consistent with relieving a generalized molecular stressor rather than activating particular therapeutic pathways.

### Clinical Significance and Sex-Specific Findings

The magnitude of hemodynamic improvement, while modest at 8-12% RVSP reduction, compares favorably to approved PH therapies and occurred alongside substantial improvements in diastolic function that are often more clinically relevant than absolute pressure reductions. Importantly, these effects were most pronounced in metabolically-driven disease models, suggesting that patient selection strategies focusing on metabolic phenotypes might optimize therapeutic response in clinical translation.

We found pronounced sex differences in both cardiac lipid metabolism and treatment response. Male right ventricles contained approximately 4-fold higher long-chain fatty acid levels compared to females (**Figure 5D**). Oddly, this may be the first report of this sex difference-we could not find other publications directly measuring LCFA in the RV, despite substantial prior work showing that sex affects RV function in PH. What makes this finding more intriguing is its ventricular specificity: the left ventricle showed no significant sex-related differences in fatty acid content. This, combined with the distinct molecular responses we observed between RV and LV even within the same model, indicates real differences in how these chambers respond to metabolic stress and lipid scavenging therapy.

These sex-specific metabolic differences may explain differential treatment responses and have important implications for clinical trial design and personalized therapy approaches. The findings underscore the critical need for adequately powered sex-stratified analyses in cardiovascular research rather than treating sex as a simple covariate.

### Ventricular Crosstalk and Molecular Insights

The molecular changes induced by 2-HOBA in the left ventricle were both distinct from those in the right ventricle within each model and varied between models. While the strong LV changes in the AKR-HFD and LNAME-HFD models are not surprising—given that the LV was also exposed to metabolic stress from the high-fat diet, the significant molecular changes observed in the LV of the PAB model were unexpected. Since the pulmonary artery band should have directly affected only the RV, the magnitude of LV molecular alterations implies substantial physiological or signaling crosstalk between ventricles that warrants further investigation.

### Study Limitations

There are several important limitations, many of which became more apparent in retrospect. Because we did not originally intent to look at sex differences-this only arose during analysis-the number of animals per group broken out by sex are less than optimal. In retrospect, single-cell RNA sequencing would have been more useful in interpreting the molecular changes. Many of the changes we found were driven by non-cardiomyocyte cell types, and a more detailed understanding of how scavenging lipids impact each cell type would have increased our ability to interpret these results. We only examined lung gene expression in the AKR-HFD model, not the LNAME-HFD model.

The AKR-HFD model, while useful for metabolic studies, has confounding strain-specific pathologies. The sequential study design, while allowing iterative refinement, resulted in some inconsistencies in measurements across models. Most critically, the molecular data about the effects of 2-HOBA across model systems and sexes are primarily for discovery—to understand what changes associate with the improvements in physiology and energy production. We do not actually know which changes are driving the improvements in physiology at this point.

### Future Directions and Clinical Translation

These findings support that reactive dicarbonyl lipids represent a common nodal pathway that is therapeutically targetable by 2-HOBA. The existing human safety data for 2-HOBA, including recent studies in atrial fibrillation patients^23, 24, 34^ and its current over-the-counter availability, supports the feasibility of clinical translation. However, several investigations would strengthen the path forward. Formal dose-finding studies in PH-relevant models are needed. Direct measurement of tissue reactive lipid levels would confirm our proposed mechanism. Combination studies with standard PH therapies should be a priority, as the modest effect sizes—while statistically significant and potentially clinically meaningful—suggest combination approaches may be needed for optimal benefit.

The pronounced sex-specific findings warrant dedicated investigation with adequate statistical power for stratified analyses. Single-cell RNA sequencing studies would help identify which molecular changes drive physiological improvements. A better understanding of which molecular outcomes are causal versus bystanders would be useful in collecting and interpreting data from clinical trials.

Clinical development should focus on PH patients with metabolic comorbidities, where we expect the greatest benefit. Biomarker development to identify optimal candidates— such as circulating ceramide levels or metabolic imaging parameters—could enhance trial efficiency. The metabolically driven nature of the most pronounced effects, combined with broader cardiovascular benefits beyond PH-specific vasodilation, suggests that 2-HOBA may be most appropriate as adjunctive rather than monotherapy.

In conclusion, this study provides evidence supporting our hypothesis that reactive lipid scavenging acts through preventing non-specific protein adduction rather than targeting particular pathways. The intervention improves both pulmonary vascular resistance and cardiac diastolic function across mechanistically distinct disease models, with effects most pronounced in metabolically driven disease. The model-specific molecular responses despite consistent physiological benefit, combined with improvements across multiple biological systems, support a systems-level approach to oxidative stress.

The novel mechanism of action, existing human safety data, and potential for combination therapy support clinical development of this approach. The sex-specific metabolic insights—particularly the striking 4-fold difference in RV fatty acid content between males and females—provide guidance for patient selection and trial design. While questions remain about which specific molecular changes drive the observed physiological improvements, scavenging of reactive lipids may both reduce pulmonary vascular resistance and increase cardiac function in human PH as well. The breadth of beneficial effects and safety profile warrant advancement to carefully designed human studies in metabolically phenotyped PH populations.

---

**Supplemental Figure 1:**
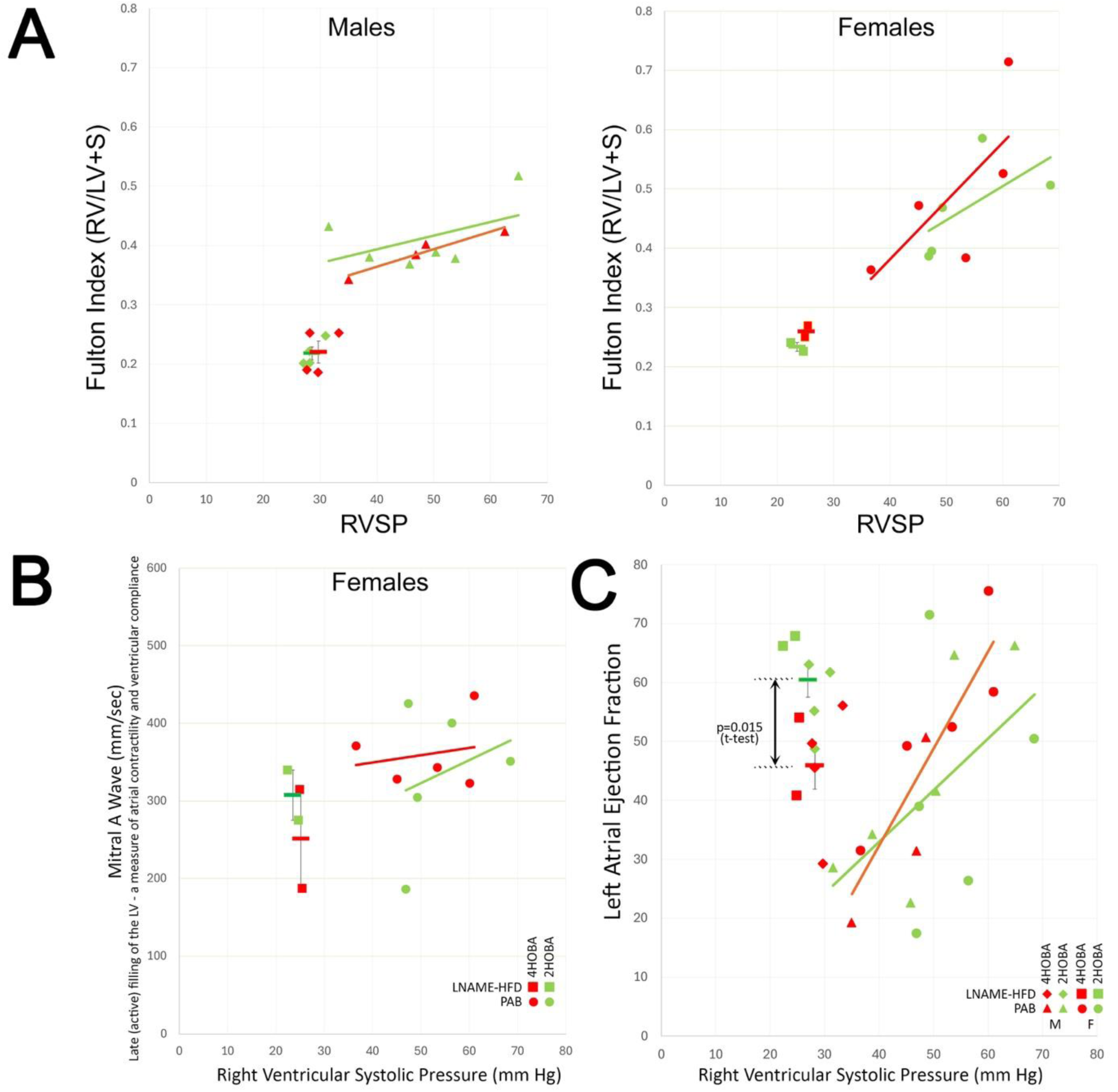
Hemodynamics. **(A)** Fulton index correlates to RVSP, but is not affected by 2-HOBA treatment.**(B)** Unlike in males, Mitral A Wave is not changed by 2-HOBA in females in LHFD and PAB models. **(C)** Left atrial ejection fraction is increased by 2-HOBA in the LNAME-HFD model, but not in PAB, where it correlates with RVSP.

**Supplemental Figure 2:**
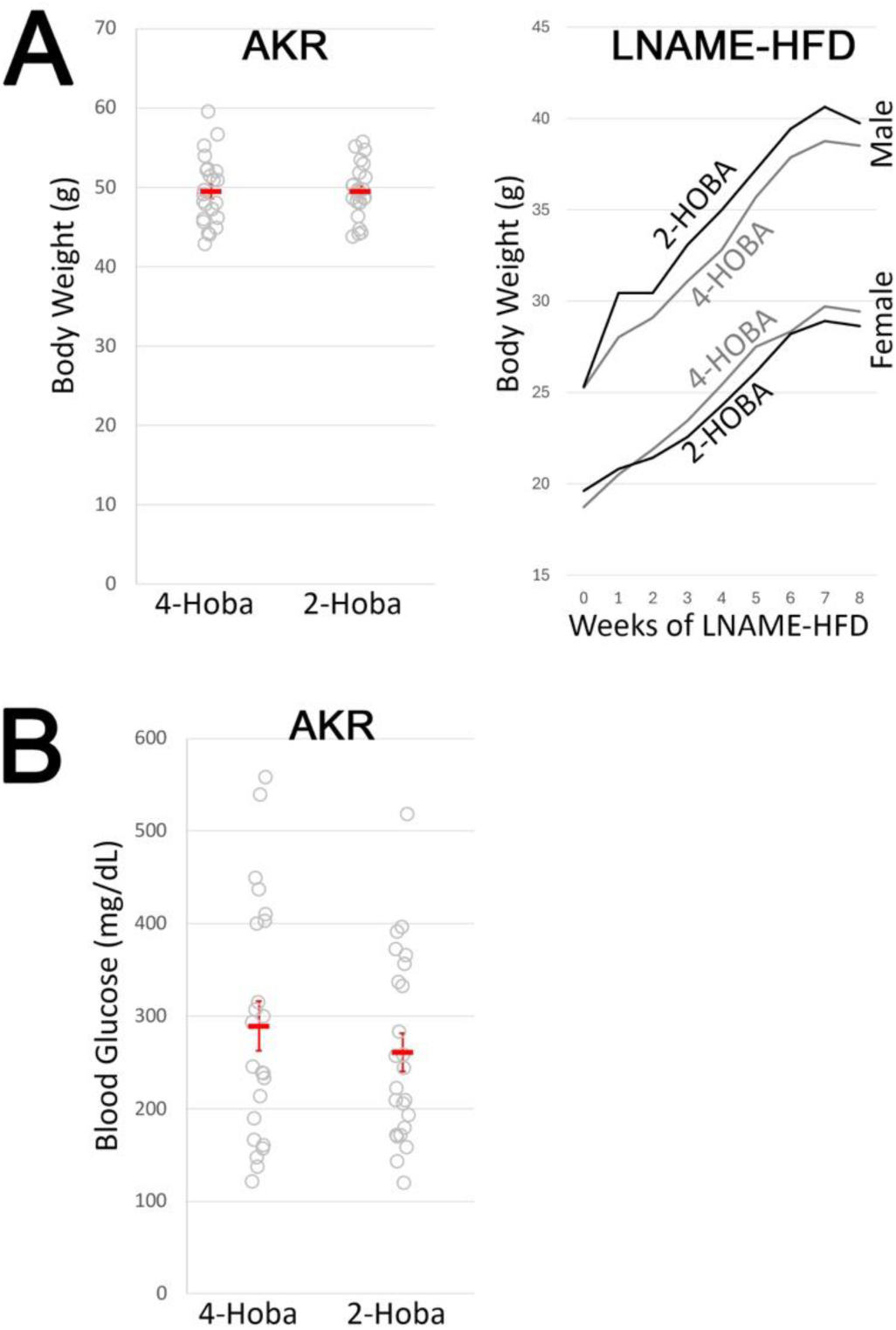
**(A)** Body weight is and rate of weight gain is not affected by 2-HOBA in AKR or LNAME-HFD models.**(B)** Blood glucose is not affected by 2-HOBA in the AKR-HFD model (not measured in other models).

## References

1. Egnatchik RA, Brittain EL, Shah AT, Fares WH, Ford HJ, Monahan K, Kang CJ, Kocurek EG, Zhu S, Luong T, Nguyen TT, Hysinger E, Austin ED, Skala MC, Young JD, Roberts LJ, 2nd, Hemnes AR, West J and Fessel JP. Dysfunctional BMPR2 signaling drives an abnormal endothelial requirement for glutamine in pulmonary arterial hypertension. Pulm Circ. 2017;7:186–199.

2. Hansmann G, Wagner RA, Schellong S, Perez VA, Urashima T, Wang L, Sheikh AY, Suen RS, Stewart DJ and Rabinovitch M. Pulmonary arterial hypertension is linked to insulin resistance and reversed by peroxisome proliferator-activated receptor-gamma activation. Circulation. 2007;115:1275–84.

3. Lane K, Talati M, Austin E, Hemnes A, Johnson J, Fessel J, Blackwell T, Mernaugh R, Robinson L, Fike C, Roberts Ii L and West J. Oxidative injury is a common consequence of BMPR2 mutations. Pulmonary Circulation. 2011;1:72–83.

4. Michelakis ED, Gurtu V, Webster L, Barnes G, Watson G, Howard L, Cupitt J, Paterson I, Thompson RB, Chow K, O’Regan DP, Zhao L, Wharton J, Kiely DG, Kinnaird A, Boukouris AE, White C, Nagendran J, Freed DH, Wort SJ, Gibbs JSR and Wilkins MR. Inhibition of pyruvate dehydrogenase kinase improves pulmonary arterial hypertension in genetically susceptible patients. Sci Transl Med. 2017;9.

5. Pugh ME, Robbins IM, Rice TW, West J, Newman JH and Hemnes AR. Unrecognized glucose intolerance is common in pulmonary arterial hypertension. J Heart Lung Transplant. 2011;30:904–11.

6. Zamanian RT, Hansmann G, Snook S, Lilienfeld D, Rappaport KM, Reaven GM, Rabinovitch M and Doyle RL. Insulin resistance in pulmonary arterial hypertension. Eur Respir J. 2009;33:318–24.

7. Talati M and Hemnes A. Fatty acid metabolism in pulmonary arterial hypertension: role in right ventricular dysfunction and hypertrophy. Pulm Circ. 2015;5:269–78.

8. Brittain EL, Talati M, Fessel JP, Zhu H, Penner N, Calcutt MW, West JD, Funke M, Lewis GD, Gerszten RE, Hamid R, Pugh ME, Austin ED, Newman JH and Hemnes AR. Fatty Acid Metabolic Defects and Right Ventricular Lipotoxicity in Human Pulmonary Arterial Hypertension. Circulation. 2016;133:1936–44.

9. Archer SL, Fang YH, Ryan JJ and Piao L. Metabolism and bioenergetics in the right ventricle and pulmonary vasculature in pulmonary hypertension. Pulm Circ. 2013;3:144–52.

10. Assad TR and Hemnes AR. Metabolic Dysfunction in Pulmonary Arterial Hypertension. Curr Hypertens Rep. 2015;17:20.

11. Talati MH, Brittain EL, Fessel JP, Penner N, Atkinson J, Funke M, Grueter C, Jerome WG, Freeman M, Newman JH, West J and Hemnes AR. Mechanisms of Lipid Accumulation in the Bone Morphogenetic Protein Receptor Type 2 Mutant Right Ventricle. Am J Respir Crit Care Med. 2016;194:719–28.

12. Hemnes AR, Brittain EL, Trammell AW, Fessel JP, Austin ED, Penner N, Maynard KB, Gleaves L, Talati M, Absi T, Disalvo T and West J. Evidence for right ventricular lipotoxicity in heritable pulmonary arterial hypertension. Am J Respir Crit Care Med. 2014;189:325–34.

13. Sakao S, Daimon M, Voelkel NF, Miyauchi H, Jujo T, Sugiura T, Ishida K, Tanabe N, Kobayashi Y and Tatsumi K. Right ventricular sugars and fats in chronic thromboembolic pulmonary hypertension. Int J Cardiol. 2016;219:143–9.

14. Paulin R, Dromparis P, Sutendra G, Gurtu V, Zervopoulos S, Bowers L, Haromy A, Webster L, Provencher S, Bonnet S and Michelakis ED. Sirtuin 3 deficiency is associated with inhibited mitochondrial function and pulmonary arterial hypertension in rodents and humans. Cell Metab. 2014;20:827–839.

15. Tang X, Chen XF, Chen HZ and Liu DP. Mitochondrial Sirtuins in cardiometabolic diseases. Clinical science. 2017;131:2063–2078.

16. Davies SS, Brantley EJ, Voziyan PA, Amarnath V, Zagol-Ikapitte I, Boutaud O, Hudson BG, Oates JA and Roberts LJ, 2nd. Pyridoxamine analogues scavenge lipid-derived gamma-ketoaldehydes and protect against H2O2-mediated cytotoxicity. Biochemistry. 2006;45:15756–67.

17. Zagol-Ikapitte I, Amarnath V, Bala M, Roberts LJ, 2nd, Oates JA and Boutaud O. Characterization of scavengers of gamma-ketoaldehydes that do not inhibit prostaglandin biosynthesis. Chem Res Toxicol. 2010;23:240–50.

18. West J, Fagan K, Steudel W, Fouty B, Lane K, Harral J, Hoedt-Miller M, Tada Y, Ozimek J, Tuder R and Rodman DM. Pulmonary hypertension in transgenic mice expressing a dominant-negative BMPRII gene in smooth muscle. Circ Res. 2004;94:1109–14.

19. Meng Q, Lai YC, Kelly NJ, Bueno M, Baust JJ, Bachman TN, Goncharov D, Vanderpool RR, Radder JE, Hu J, Goncharova E, Morris AM, Mora AL, Shapiro SD and Gladwin MT. Development of a Mouse Model of Metabolic Syndrome, Pulmonary Hypertension, and Heart Failure with Preserved Ejection Fraction. Am J Respir Cell Mol Biol. 2017;56:497–505.

20. van Ham WB, Kessler EL, Oerlemans M, Handoko ML, Sluijter JPG, van Veen TAB, den Ruijter HM and de Jager SCA. Clinical Phenotypes of Heart Failure With Preserved Ejection Fraction to Select Preclinical Animal Models. JACC Basic Transl Sci. 2022;7:844–857.

21. Agrawal V, Kropski JA, Gokey JJ, Kobeck E, Murphy MB, Murray KT, Fortune NL, Moore CS, Meoli DF, Monahan K, Ru Su Y, Blackwell T, Gupta DK, Talati MH, Gladson S, Carrier EJ, West JD and Hemnes AR. Myeloid Cell Derived IL1beta Contributes to Pulmonary Hypertension in HFpEF. Circ Res. 2023;133:885–898.

22. Townsend LK and Steinberg GR. AMPK and the Endocrine Control of Metabolism. Endocr Rev. 2023;44:910–933.

23. Pitchford LM, Driver PM, Fuller JC, Jr., Akers WS, Abumrad NN, Amarnath V, Milne GL, Chen SC, Ye F, Roberts LJ, 2nd, Shoemaker MB, Oates JA, Rathmacher JA and Boutaud O. Safety, tolerability, and pharmacokinetics of repeated oral doses of 2-hydroxybenzylamine acetate in healthy volunteers: a double-blind, randomized, placebo-controlled clinical trial. BMC Pharmacol Toxicol. 2020;21:3.

24. Pitchford LM, Rathmacher JA, Fuller JC, Jr., Daniels JS, Morrison RD, Akers WS, Abumrad NN, Amarnath V, Currey PM, Roberts LJ, Oates JA and Boutaud O. First-in-human study assessing safety, tolerability, and pharmacokinetics of 2-hydroxybenzylamine acetate, a selective dicarbonyl electrophile scavenger, in healthy volunteers. BMC Pharmacol Toxicol. 2019;20:1.

25. Bowers E, Entrup GP, Islam M, Mohan R, Lerner A, Mancuso P, Moore BB and Singer K. High fat diet feeding impairs neutrophil phagocytosis, bacterial killing, and neutrophil-induced hematopoietic regeneration. J Immunol. 2025;214:680–693.

26. Sobhy MA, Bralic A, Raducanu VS, Takahashi M, Tehseen M, Rashid F, Zaher MS and Hamdan SM. Resolution of the Holliday junction recombination intermediate by human GEN1 at the single-molecule level. Nucleic Acids Res. 2019;47:1935–1949.

27. Plecita-Hlavata L, Brazdova A, Krivonoskova M, Hu CJ, Phang T, Tauber J, Li M, Zhang H, Hoetzenecker K, Crnkovic S, Kwapiszewska G and Stenmark KR. Microenvironmental regulation of T-cells in pulmonary hypertension. Front Immunol. 2023;14:1223122.

28. Huertas A, Phan C, Bordenave J, Tu L, Thuillet R, Le Hiress M, Avouac J, Tamura Y, Allanore Y, Jovan R, Sitbon O, Guignabert C and Humbert M. Regulatory T Cell Dysfunction in Idiopathic, Heritable and Connective Tissue-Associated Pulmonary Arterial Hypertension. Chest. 2016;149:1482–93.

29. Sharma S and Aldred MA. DNA Damage and Repair in Pulmonary Arterial Hypertension. Genes (Basel). 2020;11.

30. Lemaitre RN, Jensen PN, Hoofnagle A, McKnight B, Fretts AM, King IB, Siscovick DS, Psaty BM, Heckbert SR, Mozaffarian D and Sotoodehnia N. Plasma Ceramides and Sphingomyelins in Relation to Heart Failure Risk. Circ Heart Fail. 2019;12:e005708.

31. Ruozi G, Bortolotti F, Mura A, Tomczyk M, Falcione A, Martinelli V, Vodret S, Braga L, Dal Ferro M, Cannata A, Zentilin L, Sinagra G, Zacchigna S and Giacca M. Cardioprotective factors against myocardial infarction selected in vivo from an AAV secretome library. Sci Transl Med. 2022;14:eabo0699.

32. Zhang Y, Wang D, Zhao Z, Liu L, Xia G, Ye T, Chen Y, Xu C, Jin X and Shen C. Nephronectin promotes cardiac repair post myocardial infarction via activating EGFR/JAK2/STAT3 pathway. Int J Med Sci. 2022;19:878–892.

33. Dufault VM, Oestreich AJ, Vroman BT and Karnitz LM. Identification and characterization of RAD9B, a paralog of the RAD9 checkpoint gene. Genomics. 2003;82:644–51.

34. Yoneda ZT, O’Neill M, Crawford DM, Ao M, Sun L, El-Harasis MA, Pitchford L, Rathmacher JA, Montgomery J, Shen ST, Estrada JC, Saavedra PJ, Ellis CR, Richardson T, Kangasundram A, Crossley GH, Akers WS, Ye F, Roden DM, Michaud GF and Shoemaker MB. 2-Hydroxybenzylamine for Treatment of Atrial Fibrillation: A First-in-Human Clinical Pilot Trial. Circ Arrhythm Electrophysiol. 2025;18:e013378.

